# Beyond protein functions: evaluating completeness, coherence, and consistency of genome-scale function annotations

**DOI:** 10.1101/2025.07.14.664848

**Authors:** Rund Tawfiq, Maxat Kulmanov, Robert Hoehndorf

## Abstract

Protein function annotation has traditionally followed a reductionist approach, assigning functions to individual proteins acting in isolation. This paradigm treats each annotation as an independent fact, disconnected from the broader biological system. However, proteins operate within integrated cellular networks where their functions depend on genomic context and the presence of interacting partners. Here, we develop a genome-scale evaluation framework that assesses whether annotated protein functions could plausibly coexist within a living organism. We formalize three criteria grounded in systems biology principles: completeness (presence of essential functions), coherence (satisfaction of functional dependencies), and consistency (absence of mutually exclusive functions). Applying this framework to bacterial genomes, we evaluated manually curated annotations from six model organisms and computational predictions from six methods. While model organism annotations largely satisfied our constraints — with violations primarily reflecting host– pathogen interactions — all computational prediction methods systematically failed to produce biologically plausible genome-scale annotations. Methods achieved high accuracy for individual proteins yet produced incomplete metabolic pathways, incoherent protein complexes, and taxonomically impossible function combinations. These results reveal a fundamental disconnect between the reductionist annotation model and the systems-level requirements of biological organisms. Current computational methods amplify this disconnect as they are optimized for protein-level accuracy while ignoring genome-scale constraints. Our framework provides quantitative metrics for evaluating biological plausibility and establishes a foundation for developing system-aware annotation approaches. The shift from reductionist to systems-level perspectives will be essential for annotating the rapidly growing collection of sequenced genomes and metagenomes.

**Significance Statement:** Protein function prediction methods are evaluated by their accuracy on individual proteins, but proteins operate within integrated biological systems with strict functional requirements. We developed a framework that evaluates whether predicted protein functions could plausibly coexist in a living organism by checking for completeness of essential functions, coherence of functional dependencies, and consistency with biological constraints. While manually curated annotations largely satisfy these requirements, all computational prediction methods systematically fail to produce biologically viable genome-scale annotations. This reveals a fundamental disconnect between current evaluation paradigms and the systems-level requirements of biology, highlighting the need for prediction methods that consider genome-scale constraints rather than optimizing for individual protein accuracy.

---

**W**ith the rapid accumulation of sequenced genomes, protein function annotation and prediction have become important aspects of computational biology. To capture the diversity of biological functions, frameworks such as the Gene Ontology (GO) (1) have been adopted to describe and compare functions. GO provides a comprehensive, hierarchical structure for categorizing the functions of individual proteins. The standardized description of protein functions has enabled automated, large-scale comparisons within and between organisms (2).

Experimental characterization of protein functions does not scale with the amount of protein sequences becoming available. Therefore, computational methods for function prediction have been developed using many different approaches (3). These approaches can be broadly categorized by the biological principles they aim to exploit. Methods based on sequence homology, such as BLAST2GO (4) or eggNOG-mapper (5), compare novel protein sequences to databases of well-characterized proteins and assign function based on sequence similarity; methods such as InterProScan (6) match protein sequences to established collections of domain, family, and motif signatures; and machine learning approaches used multiple types of protein features and learning paradigms to predict their functions (7).

The performance of these methods is evaluated in competitions such as CAFA (Critical Assessment of Functional Annotation) (8). CAFA provides a standardized framework for assessing protein function prediction based on predefined benchmarks and evaluation methods (9). While CAFA’s standardized benchmarks have significantly advanced the field, its evaluation paradigm focuses exclusively on the accuracy of individual protein predictions, without considering whether the collective set of predictions for an organism makes biological sense.

Current annotation models, prediction tools, and evaluation frameworks share a fundamental limitation: they treat proteins as isolated entities rather than components of integrated biological systems. The annotation model used by GO, tools for protein function prediction and their evaluations focus on assigning functions to individual proteins, where the input is usually the protein’s amino acid sequence. However, the majority of functions in the GO cannot reasonably be predicted from a single protein sequence. Specifically, the GO consists of three sub-ontologies, capturing molecular functions, biological processes, and cellular components (1). Only molecular functions, which include functions such as binding to specific compounds or enzymatic activities, can be predicted from a single protein. Biological processes in the GO require multiple proteins to participate; examples are processes corresponding to biological pathways where different proteins are required to perform different steps of the pathway. For example, glycolysis requires multiple enzymes working in sequence; predicting a single glycolytic enzyme for an organism tells us nothing about whether the complete pathway can function. From a single protein *p* that is found in organism *O*, it is impossible to predict whether the process *P* occurs, even if *p*’s sequence is identical to another protein *p*^*′*^ that participates in *P* in another organism; predicting *P* is impossible because other proteins will be required to perform *P*, and information about *p* does not tell us about the presence of other proteins.

A notable exception to this protein-centric paradigm is the development of GO Causal Activity Models (GO-CAMs) (10), which explicitly represent how multiple proteins coordinate to realize biological processes. However, GO-CAMs remain limited in scope — they are primarily focused on human biology, require manual curation, and only a small number currently exist.

With the increased ability to sequence and assemble whole genomes, and meta-genomes, we can reformulate the task of protein function annotation, prediction, and evaluation and move towards proteome-scale protein function characterization: given a set of proteins (the “proteome”), the task is to describe, predict, and evaluate the functions of all proteins of the set. This reformulation enables us to ask new questions: Does the collection of predicted functions contain all essential functions for life? Are functional dependencies satisfied? Are there combinations of functions that would be impossible in living systems?

The importance of this proteome- or genome-scale perspective becomes clear when we consider that organisms are not random collections of proteins, but evolved systems with specific functional requirements and constraints. While the reductionism used in protein databases like UniProt (11) and annotation frameworks like GO (12) — where each protein is characterized in isolation — has been useful for describing our molecular understanding, it fails to capture the emergent properties that arise from protein interactions and system-level organization, and may lead to annotations that do not respect system-level properties that govern all life.

We define three fundamental properties that protein functions for a set of proteins within a proteome should have in order to allow these functions to co-exist in a biological organism. Unlike GO-CAMs, which require manual construction of specific process models, our approach leverages existing constraints already encoded in the GO structure, combined with universal biological principles that apply across all cellular life. First, ‘completeness’ ensures the presence of functions essential for life. Second, ‘consistency’ prevents the co-occurrence of mutually exclusive functions. Third, ‘coherence’ guarantees that functional dependencies are satisfied — if function *A* requires function *B*, both must be present within the proteome. Our framework specifically addresses cellular organisms — bacteria, archaea, and eukaryotes — where these constraints reflect fundamental requirements for autonomous life (13). Viruses and phages, which depend on host cellular machinery, operate under different constraints that fall outside our current scope. However, our approach may extend to microbial communities, where essential functions can be distributed across multiple species that together form a functional system.

We formalize these three properties as logical constraints that are based on the GO, thereby directly relating our notion of proteome-level “function” to existing protein-level annotations based on the GO. We apply our framework to assess bacterial genome annotations. We test whether “gold standard”, curated annotations of model organisms satisfy our constraints, and we evaluate different protein function prediction methods according to our constraints.

This application of our framework reveals a substantial contrast between manually curated annotations and computational predictions. Manually annotated model organisms largely satisfy our constraints, with violations primarily occurring when pathogen proteins are annotated with host-specific functions (e.g., wound healing). In contrast, current function prediction methods systematically fail to satisfy these constraints across all three criteria — no method we evaluated produces annotations that would represent a biologically viable organism; while function prediction methods may accurately predict individual protein functions, they fail to capture the system-level requirements of living organisms.

Our evaluation measures and results are implemented as Free Software available at https://github.com/bio-ontology-research-group/GAEF. We anticipate this framework will guide the development of next-generation function annotation methods that explicitly consider genome-scale constraints.

## Methods

### Datasets

#### Bacterial Genome Dataset

To evaluate protein function annotations, we randomly selected 1,000 complete, annotated genomes (representing approximately 0.5% of available complete bacterial genomes at the time of selection) and their corresponding protein sequences and downloaded them from the NCBI Genomes database (14) on 2024-06-01. We selected genomes from NCBI based on these criteria: assembly level “complete”; taxon: “bacteria (eubacteria)”; annotation: “annotated by NCBI RefSeq”. These genomes represent 1,000 distinct species and 556 genera, with an average of 3,197 protein-coding genes per genome.

#### Gene Ontology

Gene functions are described using the Gene Ontology (GO) (1, 12, 15). GO is a structured, controlled vocabulary that describes the functions of gene products and relationships between functions across all life. It consists of three interconnected sub-ontologies that describe different aspects of biological function: Molecular Function (MF), Biological Process (BP), and Cellular Component (CC). The Molecular Function ontology describes specific activities of gene products at the molecular level, such as enzyme catalysis, binding interactions, or transporter functions; functions in MF can be determined based on information about a single proteins. The Biological Process ontology represents functions that require multiple molecular activities working together, like cell division, immune response, or metabolic pathways; they will usually involve more than a single protein. The Cellular Component ontology characterizes the location where gene products are active within cells or in the extracellular environment, including organelles, protein complexes, and cellular structures. Here, we use all three branches of GO collectively as “function”, and “function annotation” or “function prediction” to mean the assignment of a GO class to a protein.

There are three main version of GO:

- `GO-basic`: A simplified version containing only within-ontology relationships, i.e., no axioms that cross between different sub-ontologies of GO. `GO-basic` is intended to be sufficient for most biological applications of GO, including gene set enrichment analysis (16).
- `GO`: The core version includes axioms involving multiple relation types (‘part of’, ‘regulates’, etc.) and allows relationships to cross between different sub-ontologies.
- `GO-Plus`: An extended version of GO that contains additional axioms that link to other ontologies (16), such as ChEBI (17) for chemical entities or the Celltype Ontology (18).

For our experiments, we used the `GO` and `GO-Plus` (r16-03-2025) in Web Ontology Language (OWL) (19) format. The version of GO we used includes 51,676 classes and 90,088 logical axioms, and GO-Plus includes 84,186 classes and 1,004,038 axioms, of which 297,536 are logical axioms. For our evaluation, we relied on GO-Plus axioms involving relations such as ‘has part’ (RO:0000051) and ‘occurs in’ (RO:0000066).

#### GO Function Prediction Methods

We annotated protein sequences derived from each assembly using DeepGOMeta (v1) (20), InterProScan (v5.61-93.0) (6), SPROF-GO (r05-12-2022) (21), DeepFRI (v1) (22), TALE (r03-2021) (23), and DeepGraphGO (r07-2021) (24). We selected these methods to represent diverse approaches: structural information (DeepFRI), sequence patterns (TALE, DeepGOMeta, SPROF-GO), domain signatures (InterProScan), and protein networks (DeepGraphGO).

As DeepFRI can predict functions from Protein Data Bank (PDB) (25) structure files, we mapped RefSeq (14) protein identifiers to UniProt identifiers using the mappings provided by the UniProt Knowledgebase (r02-10-2024) (11). We then used these UniProt identifiers to download protein structures from the AlphaFold Protein Structure Database (accessed 30 Oct 2024) (26). We predicted structures for proteins without available entries in the database or corresponding UniProt identifiers using ESMFold (v1.0.3) (27). We split proteins exceeding 1,024 amino acids into the minimal number of non-overlapping chunks, and used each chunk separately for structure prediction, according to the recommendations of the ESMFold developers. We then used the resulting structure (PDB format) files for the proteins in an assembly as input for DeepFRI predictions. For proteins containing multiple chunks, we concatenated the resulting annotations from all the chunks belonging to the same protein. For TALE, we split proteins exceeding 1,000 amino acids into the minimal non-overlapping number of chunks, annotated all the chunks, and concatenated the annotations of all the chunks for each protein.

Each method, except InterProScan, provides a prediction score. For evaluation, we used a prediction score threshold that maximized the protein-centric *F*_max_ measure (9) based on a time-based split of the UniProt/SwissProt database (v2023-03, r28-06–2023 to v2023-05 r08-11-2023) (see Supplementary Materials, Table S1), following the time-based evaluation procedures established by the CAFA challenge (8). We obtained a distinct threshold for each GO sub-ontology and for each method, with the exception of InterProScan which does not output a prediction score. Instead, for InterProScan, we retained all resulting annotations provided by InterProScan without any filtering. InterProScan maps to GO through curated InterPro-to-GO mappings (28).

We processed annotations resulting from all methods for downstream applications using two distinct strategies. For the first strategy, we only retained the most specific predicted GO classes for each protein by excluding more general ancestor classes. In this context, an ancestor encompasses any GO class connected through a ‘is a’, ‘part of’, and ‘occurs in’ relationship, as well as any combination thereof. This process involved traversing the GO hierarchy and eliminating any classes that were ancestors of more specific classes already associated with the protein. We followed a second strategy where we expanded the annotations by propagating each protein’s GO classes to include all ancestors in the GO hierarchy.

#### Model Organism Genome Annotations

We obtained annotations of six model organisms to test manual function annotations. We obtained annotations of *Mycobacterium tuberculosis* ATCC 25618, *Staphylococcus aureus* NCTC 8325, *Helicobacter pylori* 26695, and *Bacillus subtilis* 168 from the Gene Ontology Annotation (GOA) Database (29); *Eschericia coli* K-12 from EcoCyc (30); and *Pseudomonas aeruginosa* PAO1 from the Pseudomonas Genome Database (31) (all databases accessed 12-03-2025). We selected these six organisms as they represent well-studied bacterial models with extensive manual curation across diverse lifestyles: free-living (*E. coli, B. subtilis*), pathogenic (*S. aureus, H. pylori, M. tuberculosis*), and opportunistic (*P. aeruginosa*). We retained annotations with the following evidence codes: Experimental: `EXP, IDA, IPI, IMP, IGI, IEP`; High-throughput experimental: HTP, HDA, HMP, HGI, HEP; Computationally-derived: `ISS, ISO, ISA, IGC`; Curator or author statement: `TAS, NAS, IC`; and Electronic: `IEA`.

### Information Content

We calculated the Information Content (IC) of GO classes to measure the specificity of functional annotations based on Resnik’s formulation (32). We computed the IC of a GO class *c* for a given genome *g* as:

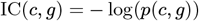

where *p*(*c, g*) is the relative frequency of class *c* in the set of all GO classes annotated to proteins in genome *g*. This formulation ensures that IC reflects the specificity of GO classes within the context of each genome, where frequently occurring (general) classes receive lower IC values, while less frequent (specific) terms receive higher IC values.

To summarize annotation specificity, we computed two metrics per genome: IC depth, defined as the mean IC across all annotations in a genome, representing the average specificity of the predicted functions; and IC breadth, defined as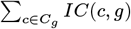 where *C*_*g*_ is the set of all GO classes annotated in *g*. To account for differences in genome size, we calculated a normalized IC breadth value by dividing the total IC by the number of proteins in the genome to capture the total functional specificity assigned per protein.

### Basic notations

To formalize our evaluation framework, we first establish the logical foundations and notation used throughout our definitions. Our aim is to extend the evaluation of protein function predictions from single proteins to entire genomes. We first introduce a set of predicates that we use in making our notions precise. Basic types (unary predicates) are *genome*(*x*), *protein*(*x*), and *function*(*x*), which allow us to distinguish between the entities we refer to. The distinction between *genome* and *protein* is introduced to reformulate the function prediction task to a task where functions of multiple proteins belonging to the same genome (or proteome) are predicted. We also use every GO class as a basic type (unary predicate), and assume that all GO classes are subclasses of *function* (i.e., for every GO class *F*, we assume ∀*x*(*F* (*x*) → *function*(*x*))). Here, we do not distinguish between the different sub-ontologies of GO.

Throughout, we will refer to specific classes of proteins (e.g., the class of proteins `FOXP2_HUMAN` with UniProt identifier O15409) as individuals, not classes; the main reason is that, here, we do not assert further axioms that pertain to proteins, and therefore can use the protein class as a logical individual. This use can also be justified ontologically and integrated with other treatments of proteins in biomedical ontologies (33). We use the same approach for genomes. Functions in GO are classes, and we will instantiate the functions when we assert that a protein has the function; proteins will have (individual) functions that are instances of some GO class. For example, to assert that the human FOXP2 protein is involved in transcriptional regulation (`GO:0006355`), we assert *hF* (*FOXP*2_*HUMAN, c*_*tr*_) GO:0006355(*c*_*tr*_), ∧ where *c*_*tr*_ is an individual instance of the GO class *Regulation of transcription, DNA-templated* (`GO:0006355`). This approach allows us to distinguish between the GO class (a universal concept) and specific instances of that function in particular proteins.

As binary predicates, we use *hF* (*x, y*) for the ‘has function’ relation between a protein *x* and a function *y*, and the newly introduced *in*(*x, y*) relation that asserts that protein *x* belongs to genome *y*. We can relate arguments of these two relations to the basic types using two axioms:

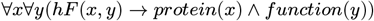

and

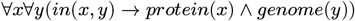

The protein function annotation, and function prediction, problem is commonly formulated as the task of assigning a set of functions (from a collection of protein functions 𝔽) to a protein *p* from a collection of proteins ℙ, i.e., to find (or learn) a function *f*_*hF*_ such that *f*_*hF*_ : ℙ → 𝒫 (𝔽) where 𝒫 *≀⊒* (𝔽) indicates the powerset of 𝔽, i.e., the set of all subsets of 𝔽. The protein *p* ∈ ℙ for which functions are predicted is given by an amino acid sequence, structure, or some other features, and the collection of functions 𝔽 is taken from an ontology such as GO, but could also refer to another vocabulary or knowledge base, such as EC numbers (enzyme prediction) (34), pathways from KEGG (35), functional domain annotations from InterPro (36) or Pfam (37), or antibiotic resistance annotations (AMR predictions) (38).

This traditional formulation treats proteins as isolated entities; however, proteins exist within the context of genomes where functional and evolutionary constraints can provide additional information and context. If whole genomes (or proteomes) are available and the relation between proteins and the genomes that code for them (as in our *in* relation), the protein function annotation or prediction task can be formulated differently: given the set of genomes 𝔾, the set of all proteins ℙ, and a set of protein functions 𝔽, find or learn a function *g*_*hF*_ such that *g*_*hF*_ : ℙ × 𝔾 *→* 𝒫 *≀⊒* (𝔽). The tuples ℙ × 𝔾 are captured by our *in* relation. This genome-centric reformulation enables us to leverage contextual information about the genomic environment in which proteins operate (the cell or organism), potentially improving function prediction accuracy by incorporating evolutionary constraints and functional relationships that exist at the genome level.

### Completeness

With the formulation of the genome-scale protein function prediction problem, we can define a set of evaluation criteria that are either defined using genome-scale function predictions directly, or generalize protein-level criteria. Our first evaluation criterion is “completeness”; the completeness condition aims to capture the idea that certain functions are necessary for all life, and one or more proteins involved in these essential functions *must* be present in every genome.

#### Definition 1

(Completeness). *Let g be a genome, let p be a protein and F is a required function from a set of required functions* ℛ *that are deemed essential for viability. A genome annotation is* complete *with respect to* ℛ *if and only if for every genome g and required function F* ∈ ℛ, *there exists at least one protein p that is in g and that has function F*. *Formally, we assert for every required function F* ∈ ℛ:

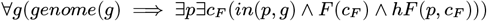

The definition captures the basic notion of completeness. While certain essential functions are shared by all life (reproduction, energy metabolism, DNA replication, etc.), some functions are only shared by certain types of organisms (e.g., only plants, or only mammals). Our notion of completeness can also be adapted to incorporate taxon-level essential functions.

While this formal definition establishes the theoretical foundation, practical implementation requires identifying which functions should be considered essential. To identify essential GO classes, we manually mapped functional categories from the *Mycoplasma mycoides* genome in Syn1.0 (39) to their corresponding GO classes (Supplementary Materials, Table S2). The Syn1.0 genome, created through systematic transposon mutagenesis experiments, represents a minimal bacterial genome required for viability under laboratory conditions. It is useful for defining essential functions as it represents an empirically validated minimal gene set. We grouped the resulting GO classes into five broad categories: Core, Defense, Glucose Metabolism, Nutrient Uptake, and Environmental Adaptation. The ‘Core’ category includes functions that are universally essential in bacteria, while the remaining categories represent functions that vary depending on metabolic strategies, environmental conditions, and stress responses. The curated set of ‘Core’ essential functions formed the basis for the target GO classes which we used to quantify the presence of essential functions in genomes.

To quantify the presence of these essential GO classes in function prediction methods, we annotated each genome in our benchmark set with GO classes predicted by each method and expanded these annotations by including the ancestor classes according to the GO hierarchy. For each GO class *F* we marked as essential, we computed the percentage of genomes *g* where at least one protein with function *F* is present, thereby quantifying the degree to which each genome satisfies our completeness criterion.

### Coherence

In addition to completeness, there are other constraints on genome-scale function annotation that rely on interactions between different functions, i.e., where the prediction of one function necessitates that another function must be present. We can define dependencies between functions either on the level of a single protein (where the prediction of a function for a protein necessitates that the protein also has another function), or on the level of a genome (where predicting a function for one protein in the genome necessitates that there is another protein in the same genome with the dependent function). We call this property *coherence*.

#### Definition 2

(Coherence). *Let g be a genome, let P* = {*p* | *in*(*p, g*)} *be the set of proteins in g, and let* 𝒞 ⊆ 𝔽 × 𝔽 *be a set of pairs of functions (which we call* dependent functions*). We call the set hF* {(*P*) = (*p, f*) | *hF* (*p, f*) *∧ p* ∈ *P*} *of function annotations for proteins in P* genome-level coherent *with respect to* 𝒞 *if and only if the following condition is satisfied for every pair* (*F*_1_, *F*_2_) ∈ 𝒞:

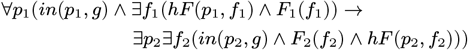

*We call the set hF* (*P*) = {(*p, f*) | *hF* (*p, f*) | *p* ∈ *P*} *of function annotations for proteins in P* protein-level coherent *with respect to* 𝒞 *if and only if the following condition is satisfied for every pair* (*F*_1_, *F*_2_) ∈ 𝒞:

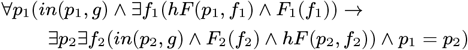

This definition captures both the notion of protein-level coherence and genome-level coherence. From the definitions, it is obvious that protein-level coherence is necessary for genome-level coherence but not the other way around.

One potential source of the set 𝒞 of dependent functions are the axioms of the GO itself; the GO axioms allow us to generate sets of dependent functions *C* both for protein-level and genome-level coherence. The GO mainly uses the “true path rule” that ensures that protein-level annotations are propagated across the GO hierarchy to ensure protein-level coherence (12); here, we focus primarily on genome-level coherence.

#### Evaluating Pathway and Process Coherence

We can use the axioms in the GO to identify genome-scale dependencies using axioms involving the ‘has part’ (`RO:0000051`) relation. We define the set of dependent functions 𝒞_*hp*_, for every GO class *F* :

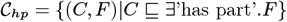

The intuition of this set of dependent functions is that there are processes that have necessary parts (or “steps”) that must be executed for the entire process to take place; however, these steps can be performed by (potentially) multiple proteins, not necessarily by a single one. This results in a genome-level coherence constraint. An example of this constraint is *Lipopolysaccharide immune receptor activity* (`GO:0001875`) which requires, through a ‘has part’ (`RO:0000051`) relation, *Lipopolysaccharide binding* (`GO:0001530`).

We used the ELK reasoner (40) to find all sub-classes of the class “has part some X” (for every class X in GO), retaining only direct sub-classes; use of ELK is necessary as we are not only interested in asserted dependencies but also those that can be inferred deductively. As a result, we obtained 5,038 pairs of classes connected through ‘has part’ relations.

To assess process coherence with respect to these pairs of classes, we calculated, at the genome level, the proportion of ‘has part’ relations that were satisfied in a genome’s annotations. Let ℛ be the set of all (*C, F*) ∈ 𝒞 _*hp*_ where *C* is present in genome *g*:

Let ℛ = {(*C, F*) ∈ 𝒞 _*hp*_ |*∃p∃ c*_*C*_(*in*(*p, g*) *∧ C*(*c*_*C*_) *∧ hF* (*p, c*_*C*_))}, and let ℳ be the subset of ℛ where the required part *F* is missing: ℳ = {(*C, F*) ∈ ℛ |*¬ ∃ p ∃ c*_*F*_ (*in*(*p, g*) *∧ F* (*c*_*F*_) *∧ hF* (*p, c*_*F*_))}. We then calculated process coherence as the percentage of satisfied dependencies:

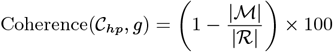

This kind of dependency can be extended to molecular pathways, which represent coordinated series of biochemical reactions (41). In metabolic pathways, these reactions transform substrates into products through enzyme-catalyzed steps, where each enzyme corresponds to a specific molecular function. Specifically, if a GO class *F*_*P*_ is mapped to a pathway 𝒫 and some protein *p* in genome *g* is predicted to have *F*_*P*_ as function, we assess whether the genome contains proteins with the necessary functions to execute the entire pathway.

Metabolic pathways can be represented as directed graphs where nodes are metabolites and edges are reactions catalyzed by specific enzymes (or other functional proteins). Many pathways contain alternative routes that can achieve the same overall transformation through different intermediate steps. This representation can be generalized to signaling pathways, where nodes may represent proteins or protein states (e.g., phosphorylated forms), and edges represent processes such as protein modifications, protein–protein interactions, or regulatory relationships. In both metabolic and signaling contexts, pathways often contain alternative mechanisms to achieve the same functional outcome.

We extracted 474 MetaCyc (42) pathway annotations and their corresponding GO classes from the annotation property assertions of GO. Using the Pathway Tools API (v28.0) (43), we retrieved each pathway’s structure, reactions, and sub-pathways. We recursively expanded reactions to include sub-pathways to build a complete directed graph representing the pathway, where nodes are reactions and edges represent dependencies between steps (as determined by the *get-predecessors* function in Pathway Tools). We mapped each reaction to its EC number using the `ec_number` attribute from the corresponding `PFrame` object. For reactions with multiple EC numbers, we generated all possible combinations of EC number paths through the pathway to account for alternative enzymatic routes. We mapped EC numbers to GO classes using http://www.geneontology.org/external2go/ec2go (r16-03-2025), resulting in 1,097 successful mappings and 255 unmapped entries.

For each pathway 𝒫, we define a “path” as a chain of reactions connecting input to output metabolites, where each reaction is catalyzed by a specific function (enzyme, transporter, or other protein). We identified all such paths using depth first search (DFS) on the reaction graph. A path *i* is defined as a sequence of functions [*F*_1_, *F*_2_, …, *F*_*k*_], where each *F*_*j*_ corresponds to an edge (reaction) in the graph. We denote the set of all paths in pathway 𝒫 as *I*_𝒫_ = *i*_1_, *i*_2_, …, *i*_*n*_. For each path *i* ∈ *I*_𝒫_, we define ℱ_*i*_ as the set of functions required to complete that specific path. A function *F* is considered present in genome *g* if there exists at least one protein *p* in *g* that is annotated with *F*.

We then calculated the completeness of a pathway with respect to a genome by identifying the proportion of required functions in each path that are present in the genome. Because metabolic networks often contain alternative routes that achieve the same biochemical transformation, a pathway is considered coherent if at least one complete path from pathway inputs to outputs exists. Therefore, we score pathway coherence using the maximum completeness across all possible paths:

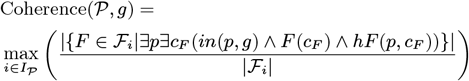

This pathway coherence measure implements the formal genome-level coherence definition by treating each pathway as a set of function dependencies 𝒞 _𝒫_,where functions in the same path are dependent on each other for the pathway’s operation. Instead of a binary determination of coherence, this metric quantifies the degree to which the coherence condition is satisfied for alternative paths through the pathway.

#### Evaluating Protein Complexes

Protein complexes require the presence of multiple interacting proteins. Homomeric complexes can form from multiple copies of the same type of protein, and heteromeric complexes require multiple distinct types of proteins. We evaluated whether heteromeric protein complex annotations are coherent so that, if a protein *p* in genome *g* is annotated with a class representing a heteromeric protein complex, there is at least another protein *p*^*′*^ in *g* that has the same annotation.

We evaluated the coherence of protein–protein complexes in bacterial genomes using GO annotations. First, we identified protein complex classes (children of *Protein–protein complex*, `GO:0032991`). We manually reviewed its subclasses and classified them as homomers (complexes of identical protein subunits), heteromers (complexes of distinct protein subunits), or classes that could represent either type. We used this curated set (available on Github https://github.com/bio-ontology-research-group/GAEF) to identify the type of protein–protein complex.

For each genome, we extracted all proteins annotated with *Protein–protein complex* (`GO:0032991`) or any of its child classes. We then classified these annotations into two categories: (1) coherent annotations, which include annotations of homomeric complex classes to any number of proteins, and annotations of heteromeric complex classes to multiple (at least two) different proteins within the same genome; (2) incoherent annotations, which include annotations of heteromeric complex classes to exactly one protein within the genome. For a genome *g* with a set of proteins *P* = {*p*| *in*(*p, g*)}, and for a complex class *C ⊑* GO:0032991, we define the coherence of its annotation as:

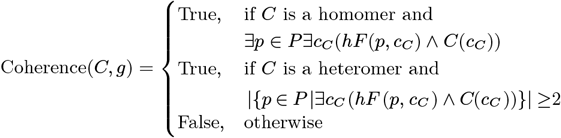

For each genome, we calculated protein complex coherence by dividing the number of incoherent complexes (heteromeric complexes annotated to exactly one protein in the genome) by the total number of complex classes annotated in the genome:

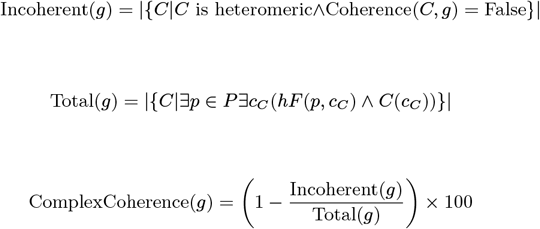

### Consistency

While coherence captures positive dependencies (required co-occurrence), consistency addresses negative dependencies (mutual exclusion). Coherence captures the notion of dependencies between functions: if there is a protein with function *F*_1_, there must be a protein with function *F*_2_. We also consider another type of dependency where functions exclude each other: having a protein with function *F*_1_ in a genome excludes having a protein with function *F*_2_ in the same genome.

#### Definition 3

(Consistency). *Let g be a genome, let P* = {*p* | *in*(*p, g*)} *be the set of proteins in g, and let* ℳ ⊆ 𝔽 × 𝔽 *be a set of pairs of functions (which we call* mutually exclusive functions*). We call the set hF* (*P*) = {(*p, f*)|*hF* (*p, f*) *∧ p* ∈ *P*} *of function annotations for proteins in P* genome-level consistent *with respect to* ℳ *if and only if the following condition is satisfied for every pair* (*F*_1_, *F*_2_) ∈ ℳ:

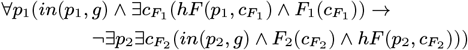

*We call the set hF* (*P*) = {(*p, f*) | *hF* (*p, f*) *∧ p* ∈ *P*} *of function annotations for proteins in P* protein-level consistent *with respect to* ℳ *if and only if the following condition is satisfied for every pair* (*F*_1_, *F*_2_) ∈ ℳ:

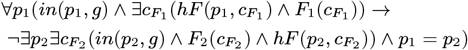

Consistency ensures that no “contradictory” functions are present in function annotations. An example would be the annotation of *photosynthesis* and *neuron development* for proteins on the same genome, or any two functions where one is restricted to multicellular organisms and the other to single-cellular organisms. The *in taxon* constraints expressed in the GO (44), where certain functions are limited to occuring only in certain taxa and others never in certain taxa, can be used to identify mutually exclusive functions and measure consistency.

To evaluate taxonomic consistency, we use the ‘in taxon’ constraints encoded in GO. The constraints specify which taxa can (‘only in taxon’, `RO:0002160`) or cannot (‘never in taxon’, `RO:0002161`) perform certain functions. We extracted the constraints from the `go-computed-taxon-constraints.obo` file (r16-03-2025), which provides curated taxonomic restrictions for GO classes (44).

We deliberately avoid comparing predicted functions against the known taxonomy of the sequenced genome for two reasons: First, taxonomic assignments of novel genomes may be uncertain. Second, in metagenomic applications, sequences may come from unknown or unculturable organisms. By checking for logical consistency among the annotated functions themselves, our framework remains applicable even without taxonomic identity.

For each genome, we aggregate all taxonomic constraints across all annotated proteins; for a set of proteins, we obtain the set of positive (‘only in taxon’) and negative (‘never in taxon’) constraints. Given a set of GO function ℱ, let *p*_1_, …, *p*_*n*_ be the set of taxa which occur in positive (‘only in taxon’) restrictions in ℱ, and *n*_1_, …, *n*_*m*_ the set of taxa with occur in negative restrictions. From the taxon constraints, we construct the logical expression *p*_1_ *⊓*… *⊓p*_*n*_*⊓¬ n*_1_ … *⊓n*_*m*_ and test whether the resulting class is satisfiable when combined with the axioms of NCBI Taxonomy (45).

Because the ELK reasoner is not able to process negated classes, for each class *C* that occurs in a negative GO taxon constraint, we create a new class *C*_*neg*_ that we declare to be disjoint with *C*; the reason for this is that ELK can process disjointness between named classes, but not direct negation. We use these newly created classes to test for satisfiability of the taxon constraints. When a contradiction is detected (i.e., the class expression *C* is unsatisfiable, *C ⊑ ⊥*), we utilize ELK’s explanation capability to generate logical justifications for the inconsistency, which can help identify the specific annotations and taxon constraints that conflict.

When evaluating model organism annotations specifically, we excluded annotations with evidence code `ISS`. Our analysis revealed that ISS annotations, which are computationally derived (using sequence or structural similarity) accounted for the majority of taxonomic consistency violations in curated databases.

### CAFA Evaluation Comparison

While our three criteria assess the biological plausibility of genome-scale annotations, it remains unclear how they relate to benchmarks focused on individual protein accuracy. To address this, we systematically compared our metrics (completeness, coherence, consistency) with CAFA’s standard performance measures, as CAFA (8) is the current gold standard for function prediction evaluation.

CAFA evaluates methods using *F*_*max*_ (maximum F-score across all thresholds) and *S*_*min*_ (minimum semantic distance) (9), both computed on held-out test proteins. These metrics capture prediction accuracy at the individual protein level, while our metrics evaluate system-level biological plausibility. We tested whether methods optimized for CAFA performance also produce complete, coherent, and consistent genome-scale annotations.

To test this hypothesis, we computed CAFA metrics for five prediction methods (DeepFRI, DeepGOMeta, DeepGraphGO, TALE, and SPROF-GO) using a time-based split of UniProt/SwissProt (training: `v2023-03, r28-06-2023`; test: `v2023-05, r08-11-2023`); see Supplementary Materials Table S1. This temporal split mimics CAFA’s evaluation strategy where methods are tested on proteins annotated after training. We did not include InterProScan in the evaluation as it does not provide confidence scores to compute threshold-dependent metrics.

For correlation analysis, we matched evaluation contexts: when comparing completeness (which uses BPO terms), we correlated against *F*_*max*_ computed specifically for BPO. Similarly, we compared protein complex coherence against CCO-specific metrics, and pathway coherence against BPO metrics (as metabolic pathways are classified under biological processes).

We computed Spearman’s rank correlation coefficient (46) to assess whether methods that perform well on CAFA benchmarks also produce annotations that satisfy our genome-scale constraints. This analysis reveals whether the CAFA evaluation paradigm adequately captures the requirements for biologically plausible proteome annotation.

## Results

We developed a formal framework for evaluating functional annotations of proteins that quantifies the biological plausibility of proteome-wide function assignments through three complementary criteria. *Completeness* evaluates whether genomes contain the minimal functional repertoire necessary for viability. *Coherence* measures functional interdependencies at three biological levels: biological processes and their component parts, metabolic pathways and their alternative routes, and protein complexes and their subunit requirements. *Consistency* enforces constraints to prevent biologically impossible function assignments. Our framework applies to both manually curated annotations and computational predictions.

To demonstrate our framework’s utility, we applied it to both established reference annotations of six bacterial model organisms and predictions from six computational methods (DeepGOMeta (20), InterProScan (6), DeepFRI (22), TALE (23), DeepGraphGO (24), SPROF-GO (21)) across 1,000 bacterial genomes. Our analysis reveals that while model organisms largely satisfy these constraints, current computational methods systematically fail to produce biologically coherent annotations.

### Model Organism Annotations

We first established baseline expectations by evaluating annotations from well-characterized model organisms. We evaluated the completeness of essential functional categories in the bacterial model organism annotations (Figure 1). All essential function classes were present in all model organism annotations, with the exception of *Chromosome segregation* (`GO:0007059`) in *P. aeruginosa* and *Redox homeostasis* (`GO:0045454`) in *H. pylori*. This near-complete coverage of essential functions reflects that we chose well-studied organisms and validates our selection of core essential functions.

**Fig. 1.**
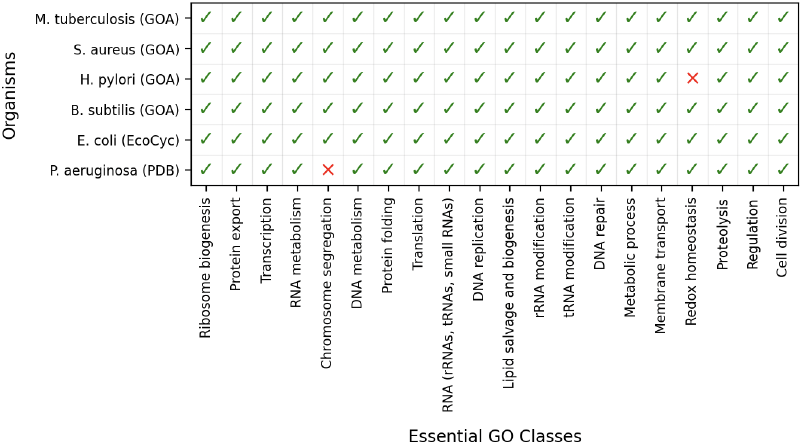
Presence or absence of ‘Core’ GO classes in genome annotations from model organisms, with checked cells indicating presence of at least one annotation to the corresponding class. Genomes were obtained from GOA, the Pseudomonas Genome Database, and EcoCyc.

Next, we evaluated the coherence of model organism annotations using process, pathway, and complex-based dependencies (Table 1). These measurements reveal how complete the functional knowledge encoded in reference databases is, and can highlight differences in our understanding of these model systems.

**Table 1.** Percentage of coherent ‘has part’ (RO:0000051) relations, MetaCyc pathway annotations, and consistent protein complexes across selected model organisms.

| Model Organism | has part | Pathways | Complexes |
| --- | --- | --- | --- |
| *B. subtilis* | 94.00% | 61.84% | 63.51% |
| *E. coli* | 96.25% | 82.84% | 84.23% |
| *H. pylori* | 92.77% | 45.10% | 73.21% |
| *M. tuberculosis* | 95.98% | 61.45% | 72.97% |
| *P. aeruginosa* | 83.90% | 20.83% | 72.73% |
| *S. aureus* | 93.29% | 48.39% | 65.67% |

*E. coli* consistently demonstrates the highest coherence across all three measures, reflecting that it is one of the most extensively-studied bacterial model organism (47). This pattern further indicates that coherence measurements effectively capture the completeness of our functional knowledge rather than merely annotation quality.

Process coherence shows the highest values across all organisms, while pathway and complex coherence scores are lower. One possible explanation is that ‘has part’ relationships are explicitly encoded in GO’s ontological structure whereas pathway and complex coherence information is not; therefore, curators are directly aware of the ‘has part’ dependencies when making annotations but not of the other dependencies.

Consequently, the coherence measurements serve as indicators of both annotation completeness and the implicit biological knowledge embedded in different curation approaches. *E. coli*’s superior performance across all metrics reflects intensive research, while the relatively high process coherence across all organisms demonstrates the effectiveness of GO’s axiomatic structure in maintaining functional dependencies.

We evaluated taxonomic consistency based on *in taxon* constraints expressed in the GO (44). Taxonomic consistency analysis revealed unexpected complexity in even well-curated annotations. *B. subtilis* and *S. aureus* satisfied taxonomic consistency constraints, while the rest of the evaluated model organism genomes contained functions with mutually exclusive taxonomic requirements. We identified three distinct causes of taxonomic consistency violations.

First, they may indicate potential annotation errors; for example, this may the case for *P. aeruginosa* where the protein `UniProt:PA5451` is annotated with *Pellicle* (`GO:0020039`), which has a ‘never in taxon’ *Bacteria* constraint. Although the annotation is manually curated, the referenced publication makes no mention of the pellicle or related structures, suggesting a potential problem in the annotation process and highlighting the need for automatic semantic checks (as implemented in our evaluation framework). Second, we found that taxonomic constraints in GO may be overly restrictive. In *E. coli*, the protein `UniProt:P60723` is annotated with *RNA-binding Transcription Regulator Activity* (`GO:0001070`), which has the ‘only in taxon’ *Viruses* constraint. However, the definition of this class describes a regulatory mechanism that is also known to occur in *E. coli* (48). Third, we found annotations that describe functions of pathogenic organisms within a host. For example, in *H. pylori* and *M. tuberculosis*, proteins `UniProt:O25743` and `UniProt:P9WQP1` are annotated with *Negative regulation of T cell proliferation* (GO:0042130) (49) and *Positive regulation of plasminogen activation* (`GO:0010756`) (50), respectively. Both proteins are correctly annotated with these functions yet perform them only during infection, and the biological processes occur in the host, not in the organisms themselves.

Beyond structural constraints, we also examined annotation specificity by calculating the average information content (IC) for each annotation (Supplementary Materials, Figure SS1). All model organism annotations demonstrated high average IC values, with *P. aeruginosa* and *E. coli* demonstrating the highest values. We also calculated the IC breadth for each organism to quantify the total functional information assigned, on average, to each protein (Supplementary Materials, Figure SS2). *E. coli* demonstrated the highest IC breadth, and *P. aeruginosa* the lowest. The inverse relationship between average IC and IC breadth in *P. aeruginosa* versus *E. coli* suggests different annotation strategies: deep but narrow versus broad but shallow annotations. Furthermore, our results indicate that specific, detailed functional assignments can be made independently of overall coherence. This suggests that annotation specificity and systematic coherence represent complementary rather than competing aspects of annotation quality.

### Application to automated function annotation methods

Having established baseline performance expectations from manually curated model organism annotations, we next evaluated six computational methods representing distinct methodological approaches to protein function prediction. This diversity allows us to assess how different types of biological evidence contribute to genome-scale annotation coherence.

#### Essential Term Completeness

Our evaluation across 1,000 bacterial genomes reveals systematic differences in how these methodological approaches perform across the three evaluation criteria. When evaluating essential function completeness (Figure 2), structural information proves most effective: DeepFRI successfully detected all 22 ‘Core’ GO classes in 100% of genomes. This performance may reflect the direct relationship between protein structure and essential catalytic functions. In contrast, sequence-based methods showed variable performance, with TALE demonstrating the lowest coverage, particularly for *Redox Homeostasis* (GO:0045454) at 1.7% and *Cell Division* (GO:0051303) at 44.2%. Notably, *Chromosome Segregation* (GO:0007059) was systematically underpredicted across methods, with DeepGOMeta failing to annotate it entirely. This pattern mirrors gaps observed in model organism annotations, suggesting challenges in recognizing these functions across both computational and manual approaches.

**Fig. 2.**
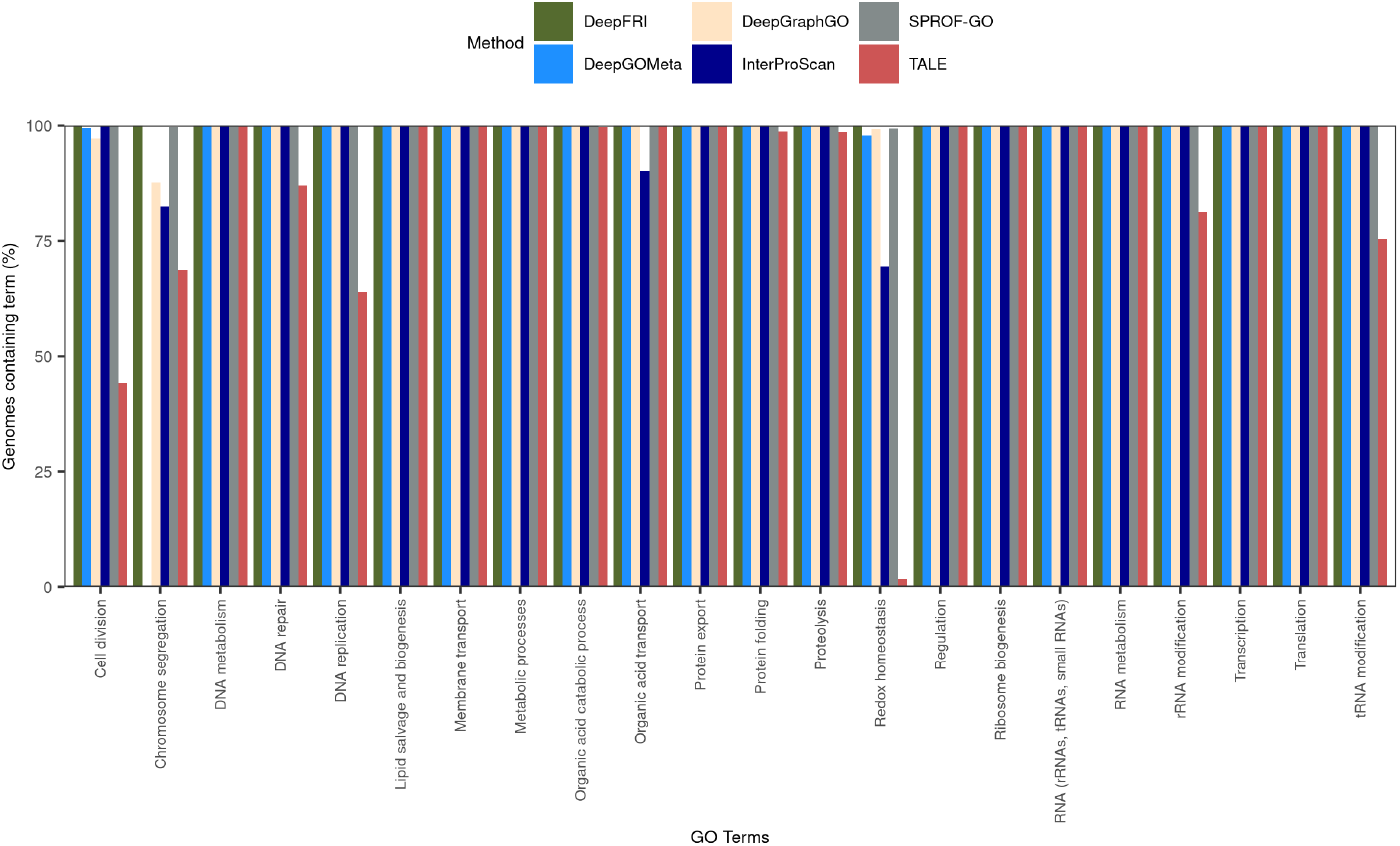
Essential GO class presence in genomes annotated using several methods. The percentage of genomes containing each ‘Core’ class by each method is shown.

#### Pathway, Process, and Protein Complex Coherence

Coherence reveals how different methodological approaches capture functional dependencies at multiple biological levels. For process coherence using ‘has part’ relations (Figure 3), methods incorporating ontological structure, protein–protein interactions, or domain knowledge (DeepGraphGO, InterProScan) achieved performance comparable to model organism annotations. DeepFRI showed the lowest process coherence, indicating that structural information alone is insufficient for capturing functional dependencies explicitly encoded in GO.

**Fig. 3.**
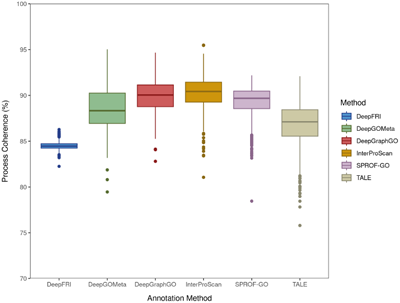
Distribution of process coherence for each method based on the percentage of missing ‘has part’ (`RO:0000051`) relations in the resulting annotations.

Pathway coherence analysis (Figure 4) demonstrated that methods based on interaction networks (DeepGraphGO) and domains (InterProScan) exceled at recovering complete metabolic pathways, achieving performance similar to curated function annotations in organisms like *H. pylori* and *S. aureus*. Sequence-based methods showed variable performance, as SPROF-GO failed to complete any pathways, while TALE performed better than other sequence-based approaches. This finding suggests that both the features and training approach influence model performance.

**Fig. 4.**
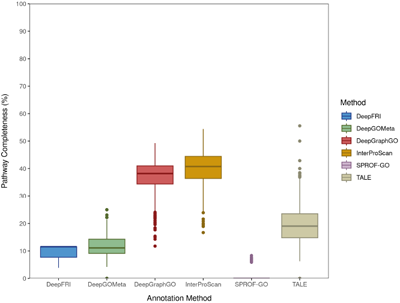
Distribution of pathway coherence for each method based on the percentage of complete MetaCyc pathways in the resulting annotations.

For protein complex coherence (Figure 5), structural information proves most valuable: DeepFRI achieved the highest coherence with minimal variation, even exceeding *E. coli* performance. This superior performance reflects the direct structural basis of protein–protein interactions in complex assembly. Surprisingly, InterProScan showed the lowest complex coherence, suggesting that domain signatures alone inadequately capture the interaction interfaces required for complex formation.

**Fig. 5.**
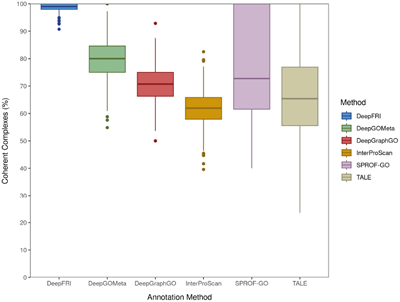
Distribution of protein complex coherence based on GO protein-protein complex annotations for each method.

#### Taxonomic Consistency

We evaluated taxonomic consistency across protein function prediction methods and identified several common patterns of constraint violations. InterProScan annoated 361 genomes as taxonomically consistent, whereas all other methods did not provide any taxonomically consistent set of genome annotations. Most methods, including TALE, InterProScan, SPROF-GO, and DeepFRI, frequently violated constraints involving virus-associated GO classes such as *Virion Component* (`GO:0044423`), *Virus Tail* (`GO:0098015`), and *Viral Capsid Assembly* (GO:0019069), which carry ‘never in taxon’ constraints for cellular organisms or bacteria.

Additionally, multiple methods showed violations involving organism-specific constraints. TALE incorrectly annotated frequently with archaeal enzymatic processes restricted to archaea, while DeepGraphGO and SPROF-GO violated constraints restricted to *Viridiplantae*, involving plant-specific cellular components and processes such as *Apoplast* (`GO:0048046`), *Plant Organ Development* (`GO:0099402`), and *Plant-type Cell Wall Organization or Biogenesis* (`GO:0071669`).

Other frequent taxonomic consistency violations involved multicellular organismal processes incompatible with bacterial genomes. DeepGOMeta primarily annotated terms related to immune processes, such as *Cytokine Production* (`GO:0001816`), while DeepGraphGO often annotated animal-specific processes like *Apoptotic process* (`GO:0006915`) which is restricted to *Opisthokonta*. DeepFRI annotations similarly violated constraints on multicellular processes such as *System Development* (`GO:0048731`).

#### Information Content

The methods we evaluated varied in annotation specificity as measured by average information content (IC) (Supplementary Materials, Figure SS3). The network-based method DeepGraphGO produced the most specific annotations on average, achieving specificity similar to *E. coli* and *P. aeruginosa*, which were the model organisms with the highest average IC. All other methods, with the exception of TALE, achieved average functional specificity comparable to other model organism annotations. This indicates that while these methods may lack coherence (i.e., they do not completely satisfy functional dependencies in the annotations they produce), they nevertheless identify very specific functional assignments.

Additionally, we evaluated total functional information per protein for each annotation method using a measure of IC breadth (Supplementary Materials, Figure SS4). DeepFRI demonstrated higher IC breadth when compared to other methods and model organism annotations. Sequence-based methods (DeepGOMeta, TALE, and SPROF-GO) also resulted in higher IC breadth than model organism annotations, while InterProScan achieved similar results, but lower than *E. coli*. These results reveal a fundamental difference between manual curation (conservative, high-confidence) and automated prediction (broad, lower-confidence) strategies.

We evaluated how some of our evaluation metrics relate to standard evaluation metrics used for CAFA across five function prediction methods (Supplementary Materials, Figure SS5). Spearman correlations revealed varying relationships between these metrics and the standard CAFA evaluation approach (*F*_max_, *S*_min_). However, no correlations were statistically significant in these comparisons based on two-sided p-values from Spearman rank correlation tests. IC depth and IC breadth did not correlate with any CAFA metric across ontologies, suggesting that prediction depth alone may not capture improvements in functional specificty.

Our results show systematic differences in how various automated function annotation approaches perform across genome-scale functional annotation tasks. Structural information shows superior performance for essential functions and protein complex relationships, while network and domain-based approaches excel at pathway-level coherence, and sequence-based methods produce highly specific annotations with variable systematic coherence. However, these performance patterns likely reflect multiple factors beyond the inherent properties of different biological information types. Specifically, the systematic performance differences we identify reveal a fundamental challenge: methods optimized for single-protein accuracy fail to capture the emergent constraints of biological systems. This gap between reductionist optimization and systems-level requirements will require new approaches that explicitly model genome-scale dependencies during training.

## Discussion

Our results demonstrate a fundamental disconnect between protein-level optimization and system-level biological requirements. While current methods effectively predict individual protein functions, they fail to produce biologically coherent genome-scale annotations. Even manually curated model organism databases showed incomplete pathway annotations and taxonomic inconsistencies, though they substantially outperformed automated methods across all three evaluation criteria — completeness, coherence, and consistency. This systematic failure across both manual and automated approaches points to deeper issues in how we conceptualize and annotate biological function.

### Limitations of the reductionist paradigm

Our evaluation revealed that even gold-standard, manually curated databases fail to capture system-level constraints. This systematic failure extends beyond methodological limitations to a fundamental issue: the GO itself embodies an inherently reductionist philosophy. Developed in the late 1990s when the first eukaryotic genomes became available (12, 51), the GO annotation model was designed to decompose biological complexity into discrete, manageable units. Each gene product receives independent functional labels, as if biology operated through isolated molecular activities rather than integrated systems.

The context-dependency of protein function reveals why genome-scale evaluation is necessary. Consider two proteins with identical sequences in different organisms: one might catalyze a step in lysine biosynthesis where all pathway enzymes are present, while the identical protein in another organism lacking the complete pathway cannot participate in this process. This example illustrates not a flaw in current annotation systems, but rather the need for formal frameworks that can express the logical implications of functional assignments. When we annotate a protein with ‘lysine biosynthesis’, we implicitly assert that other required enzymatic activities must exist somewhere in the system — but current annotation models lack the formal mechanisms to express these dependencies explicitly.

The GO was designed to capture and organize our current biological knowledge as it evolves, not to enforce system-level completeness. This design principle is valuable: it allows researchers to record what they know about individual proteins without requiring complete knowledge about how, for example, a pathway works in an organism. Annotating a protein with ‘glycolytic process’ when only some pathway enzymes are characterized is appropriate — it accurately reflects our current understanding. However, such annotations carry implicit logical commitments: they assert that other proteins with complementary functions must exist for the annotated process to occur. The challenge is designing an annotation model where we can express these logical implications.

Our coherence measure reflects the completeness of our current knowledge rather than fundamental flaws in annotation systems. When we observe incomplete metabolic pathways in well-studied organisms, this indicates gaps in functional characterization or understanding rather than errors. Similarly, when computational methods produce annotations that violate system-level constraints, this reveals their failure to capture the logical dependencies inherent in biological systems.

Recent efforts recognize this gap. GO-CAMs (Gene Ontology Causal Activity Models) (10) represent conceptual progress, explicitly modeling how multiple proteins coordinate to execute biological processes. However, they also remain limited — primarily describing human pathways through manual curation, not providing a scalable framework for genome-wide constraints. Similarly, while SBML (Systems Biology Markup Language) (52) models biochemical networks mathematically, its connection to GO remains superficial — reactions can be tagged with GO terms but without expressing functional dependencies (53). Tools like SBML Harvester (54) can detect inconsistencies between GO annotations and the model structure post-hoc, identifying when modeled reactions lack annotated cofactors, but these remain validation tools rather than integrated frameworks.

We would benefit from a formal language for biological function — one that natively expresses conditional dependencies (“This pathway functions only if enzymes *A* through *E* are present”), quantitative relationships (“Transporter expression must match metabolic flux”), emergent properties (“These five proteins together enable chemotaxis”), and mutual exclusions (“Aerobic and anaerobic respiration cannot coexist at full capacity”). Such a language would treat functions as properties of protein collectives, not individuals. Developing it requires bridging knowledge representation, systems biology, and bioinformatics; the basic logical language we use to express system-level constraints may serve as a starting point for such a language.

### Technical limitations and their effects

Beyond conceptual limitations, two technical factors explain why current methods systematically fail our evaluation criteria: First, the hierarchical structure of GO combined with CAFA’s *F*_*max*_ optimization will create a bias toward predicting general, high-confidence terms rather than specific functions. Methods maximize *F*_*max*_ by making safe predictions at higher ontology levels, avoiding the risk of incorrect specific annotations. This explains the paradox of high IC depth but poor pathway completion: methods predict broadly applicable functions while missing the specific enzymatic activities required for complete pathways. An IC-weighted *F*_*max*_ metric (9) would address this by rewarding specific predictions, but has not been widely adopted.

Second, the sequence-based data splits used in training protein function prediction models can create systematic blind spots. When methods are trained on sequence-similarity-based splits, entire protein families — and all their associated functions — may be absent from training data. This violates the assumption that training and test distributions share the same label space, leading to functions that can never be predicted regardless of model sophistication. For example, if no chromosome segregation proteins appear in training data, methods cannot learn to recognize this essential function. This explains why certain functions show 0% prediction rates across methods despite being essential for life.

## Limitations

Our framework has several limitations that should guide its interpretation and future extensions. We focused exclusively on bacterial genomes, while archaea and eukaryotes present distinct challenges through cellular compartmentalization. More fundamentally, our genome-centric approach cannot evaluate distributed metabolic systems where essential functions span multiple organisms. In syntrophic communities, “incomplete” pathways in individual genomes may represent functional partnerships where metabolites are exchanged between species.

Our essential function set derives from a single minimal genome (Syn1.0), which may not capture alternative life strategies or community-dependent organisms that rely on metabolic complementarity. The pathway analysis depends on MetaCyc coverage, missing organism-specific pathways, and assumes complete pathways exist within single genomes rather than distributed across species boundaries.

Finally, we evaluated methods that are not designed for genome-scale coherence — their low performance reflects optimization for different objectives rather than fundamental algorithmic limitations. Future work should expand to diverse taxonomic groups, develop meta-organism evaluation frameworks for microbial communities, and integrate additional pathway databases. Such extensions would require formalizing functional coherence across multiple cooperating genomes rather than single organisms.

## ACKNOWLEDGMENTS

We acknowledge support from the KAUST Supercomputing Laboratory. This work has been supported by funding from King Abdullah University of Science and Technology (KAUST) Office of Sponsored Research (OSR) under Award No. URF/1/5041-01-01, REI/1/5235-01-01, REI/1/4938-01-01, and REI/1/5659-01-01. This work was supported by funding from King Abdullah University of Science and Technology (KAUST) – KAUST Center of Excellence for Smart Health (KCSH), under award number 5932, and by funding from King Abdullah University of Science and Technology (KAUST) – Center of Excellence for Generative AI, under award number 5940.

