## Supplementary Materials for "Beyond protein functions: evaluating completeness, coherence, and consistency of genome-scale function annotations"

PNAS

**This PDF file includes:**

Figs. S1 to S7
Tables S1 to S2

### Annotation Retention Threshold

We used the confidence scores produces by each method to filter the resulting annotations for all methods, with the exception of InterProScan due to the lack of confidence scores for the resulting predictions. The threshold at which we retained the annotations is determined as the threshold that maximized $F_{\max}$ in the time-based split evaluation from our previous work on DeepGOMeta (Table S1).

**Table S1. Threshold that maximized $F_{\max}$ for each Gene Ontology (GO) sub-ontology for each method.**

| Method | MFO | CCO | BPO |
|---|---|---|---|
| SPROF-GO | 0.13 | 0.54 | 0.13 |
| DeepGOMeta | 0.27 | 0.27 | 0.11 |
| TALE | 0.28 | 0.56 | 0.15 |
| DeepFRI | 0.28 | 0.01 | 0.01 |
| DeepGraphGO | 0.33 | 0.21 | 0.30 |

### Essential Function Mappings

We manually mapped functional categories determined to be essential for the survival of *Mycoplasma mycoides* from the Syn1.0 genome to GO classes (Table S2). We were able to map most functional categories directly using exact string matches to the GO class name, definition, or description. We manually reviewed every mapping to ensure accuracy. We could not find a relevant term that directly maps to the functional category 'Transport and catabolism of nonglucose carbon sources' from Syn1.0, so we split it into 'Transport of nonglucose carbon sources' and 'Catabolism of nonglucose carbon sources', and mapped the split categories to the relevant GO classes.

**Table S2. Functional categories from syn1.0 manually mapped to the most relevant Gene Ontology (GO) class.**

| Category | Function | GO Class Name | GO Class |
|---|---|---|---|
| Core | DNA metabolism | DNA metabolic process | GO:0006259 |
| | DNA replication | DNA replication | GO:0006260 |
| | DNA repair | DNA repair | GO:0006281 |
| | Transcription | DNA-templated transcription | GO:0006351 |
| | Translation | Translation | GO:0006412 |
| | Cell division | Cell division | GO:0051301 |
| | Chromosome segregation | Chromosome segregation | GO:0007059 |
| | Ribosome biogenesis | Ribosome biogenesis | GO:0042254 |
| | Protein folding | Protein folding | GO:0006457 |
| | Protein export | Protein transport | GO:0015031 |
| | RNA metabolism | RNA metabolic process | GO:0016070 |
| | rRNA modification | rRNA modification | GO:0000154 |
| | tRNA modification | tRNA modification | GO:0006400 |
| | RNA (rRNAs, tRNAs, small RNAs) | RNA biosynthetic process | GO:0032774 |
| | Proteolysis | Proteolysis | GO:0006508 |
| | Metabolic processes | Metabolic process | GO:0008152 |
| | Membrane transport | Transmembrane transport | GO:0055085 |
| | Lipid salvage and biogenesis | Lipid metabolic process | GO:0006629 |
| | Transport of nonglucose carbon sources | Organic acid transport | GO:0015849 |
| | Catabolism of nonglucose carbon sources | Organic acid catabolic process | GO:0016054 |
| | Redox homeostasis | Cell redox homeostasis | GO:0045454 |
| | Regulation | Regulation of biological process | GO:0050789 |
| Glucose Metabolism | Glycolysis | Glycolytic process | GO:0006096 |
| | Glucose transport | Glucose transmembrane transport | GO:1904659 |
| Environmental Adaptation | Mobile elements | Transposition | GO:0032196 |
| | DNA topology | DNA conformation change | GO:0071103 |
| | Lipoprotein | Lipoprotein metabolic process | GO:0042157 |
| Nutrient Uptake | Cofactor transport and salvage | Vitamin transmembrane transporter activity | GO:0090482 |
| | Acylglycerol breakdown | Acylglycerol catabolic process | GO:0046464 |
| | Nucleotide salvage | Nucleotide salvage | GO:0043173 |
| Defense | DNA restriction | DNA restriction-modification system | GO:0009307 |
| | Efflux | Export across plasma membrane | GO:0140115 |

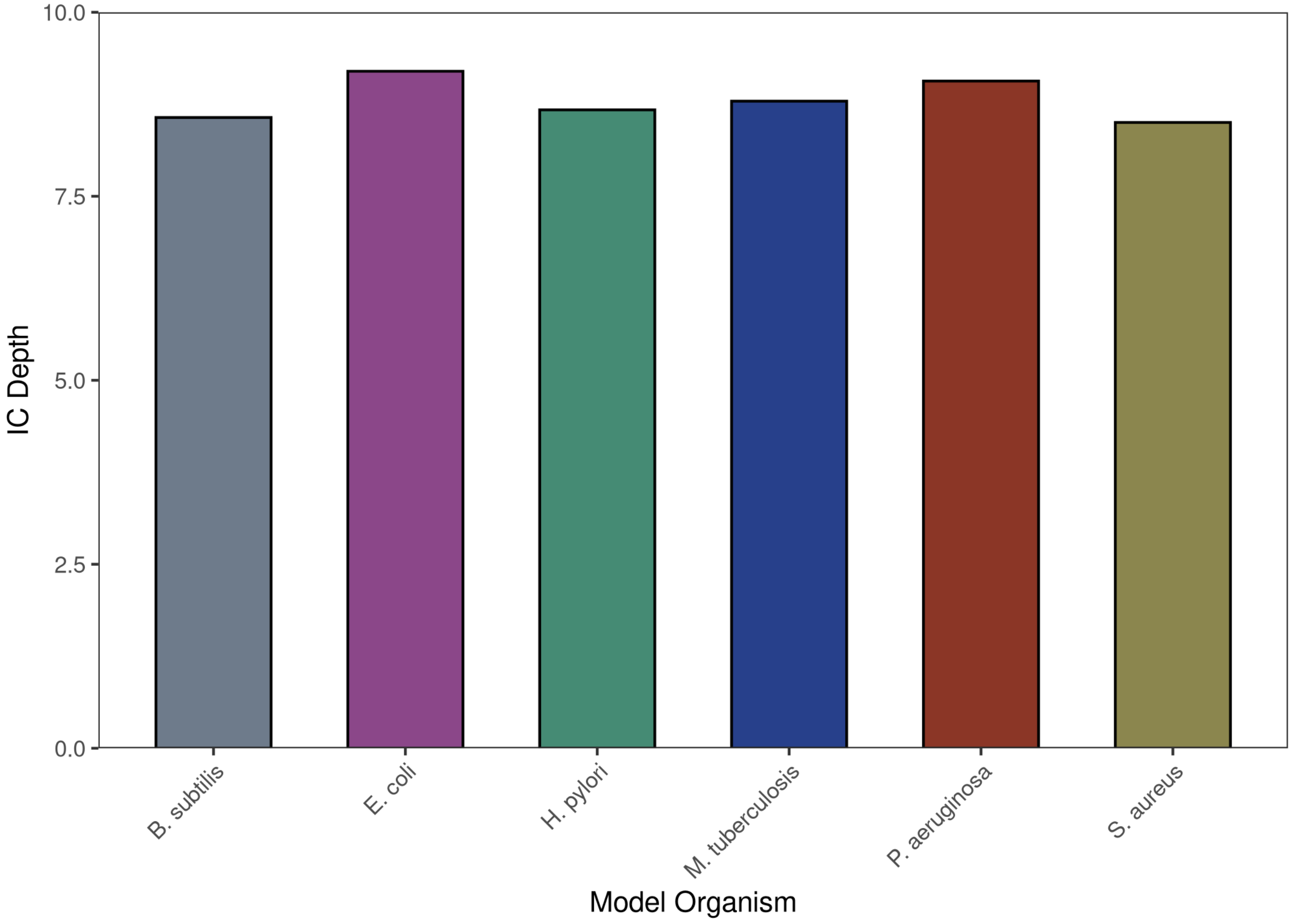


**Fig. S1.** Information Content (IC) depth for specific GO class annotations across six bacterial model organisms: *E. coli*, *B. subtilis*, *P. aeruginosa*, *H. pylori*, *S. aureus*, and *M. tuberculosis*.

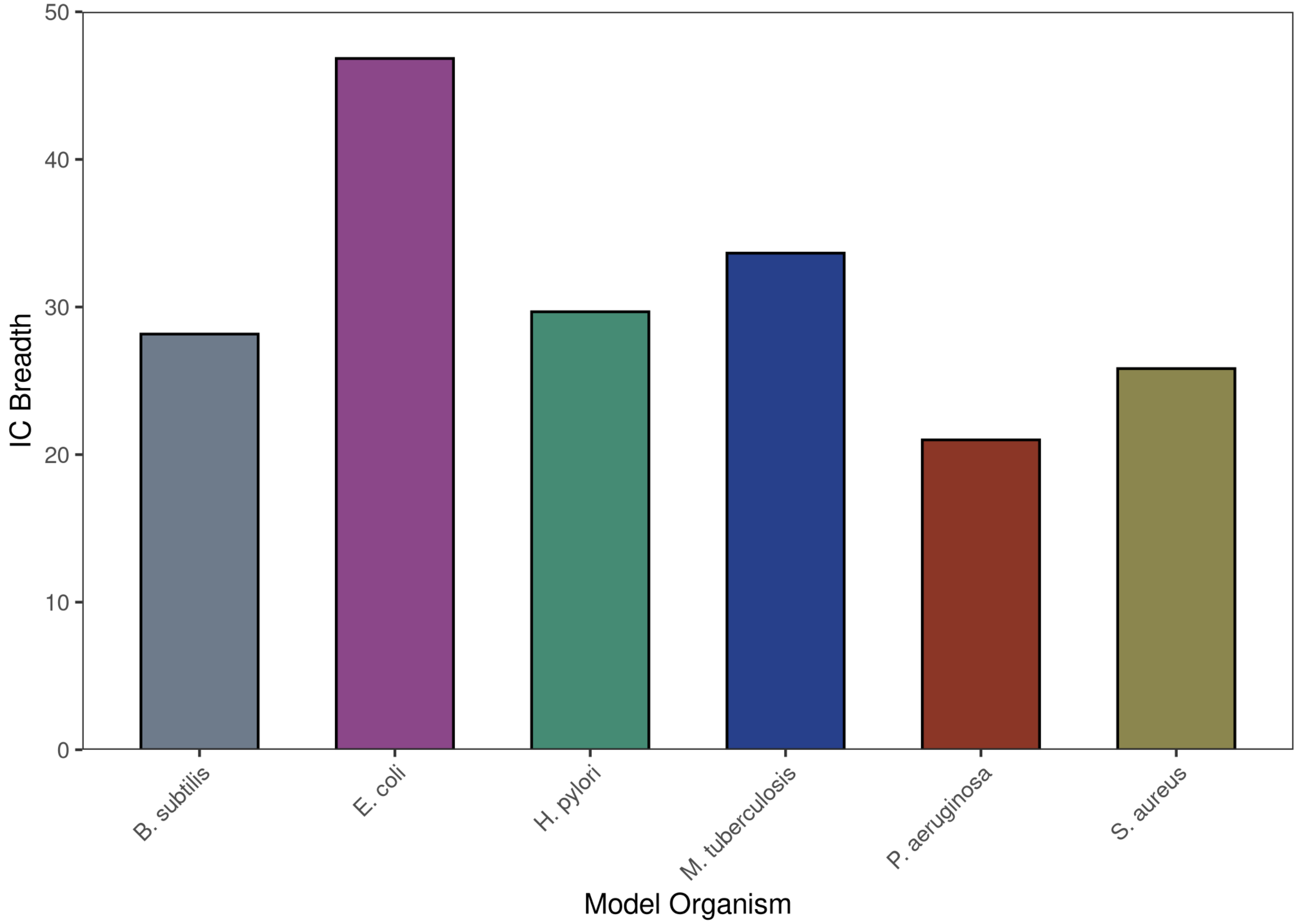


**Fig. S2.** Information Content (IC) breadth normalized by the number of proteins for specific GO class annotations across six bacterial model organisms: *E. coli*, *B. subtilis*, *P. aeruginosa*, *H. pylori*, *S. aureus*, and *M. tuberculosis*.

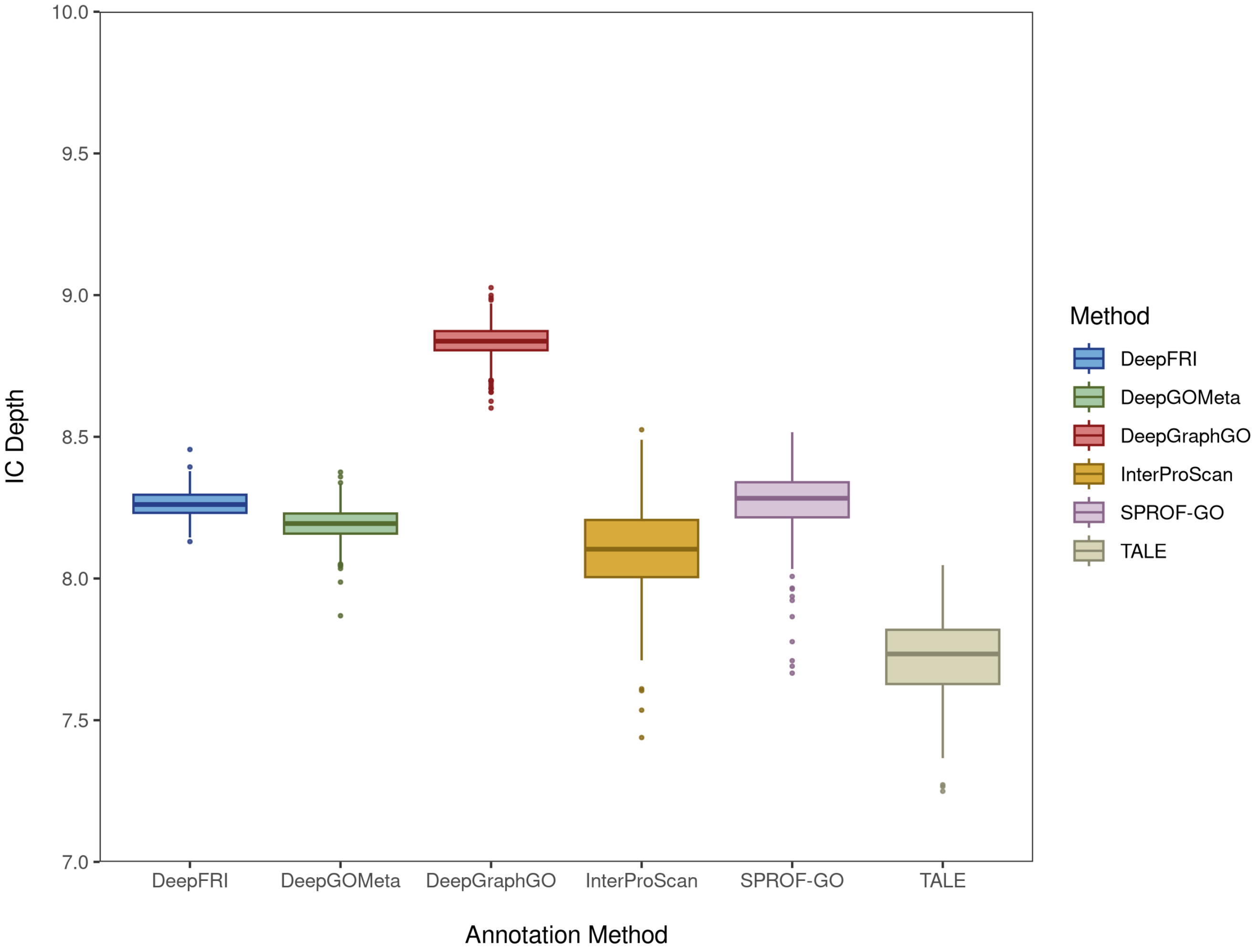


**Fig. S3.** Information Content (IC) depth for specific GO classes across six protein function prediction methods.

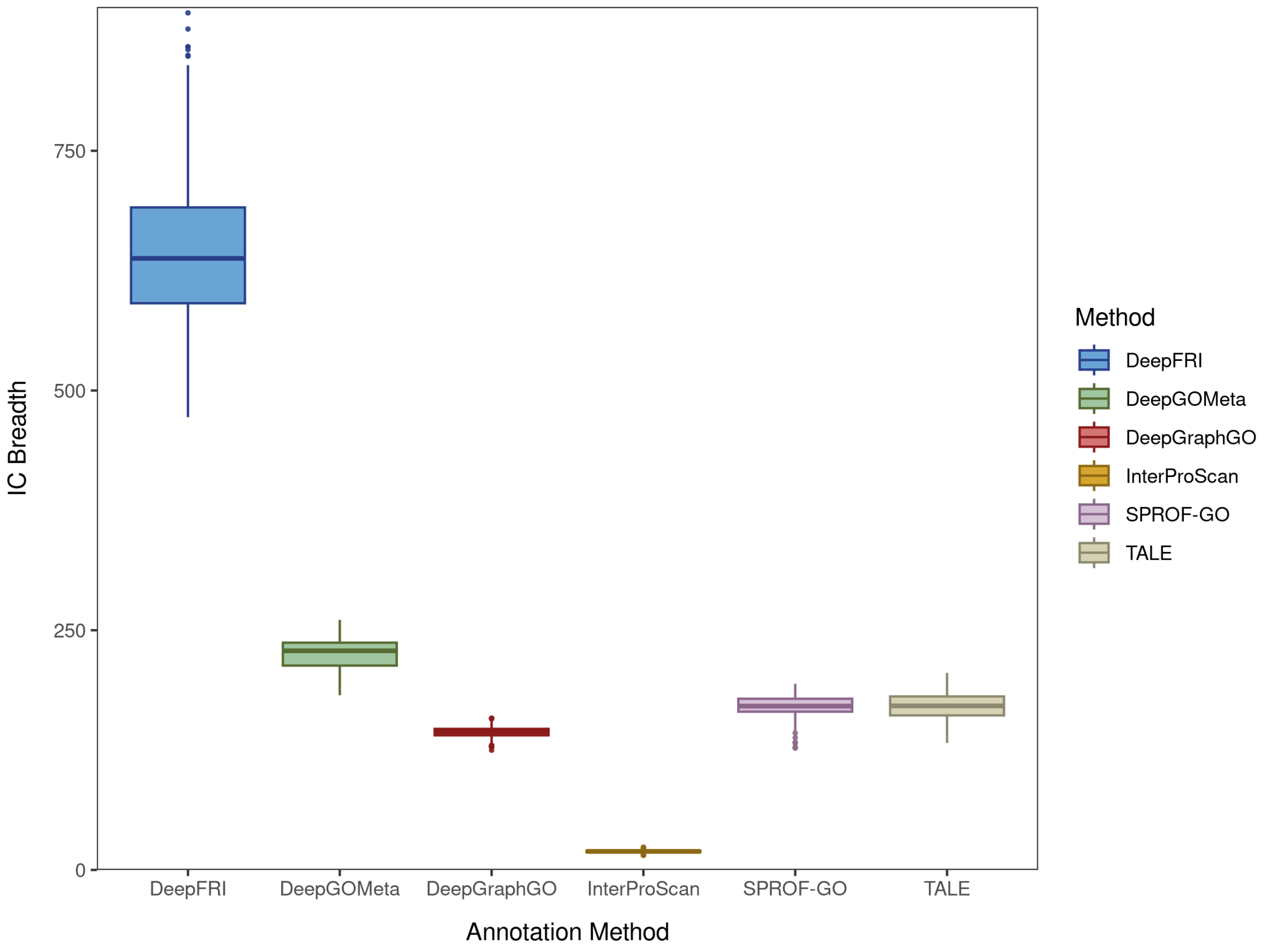


**Fig. S4.** Information Content (IC) breadth normalized by the number of proteins for specific GO classes across six protein function prediction method.

 

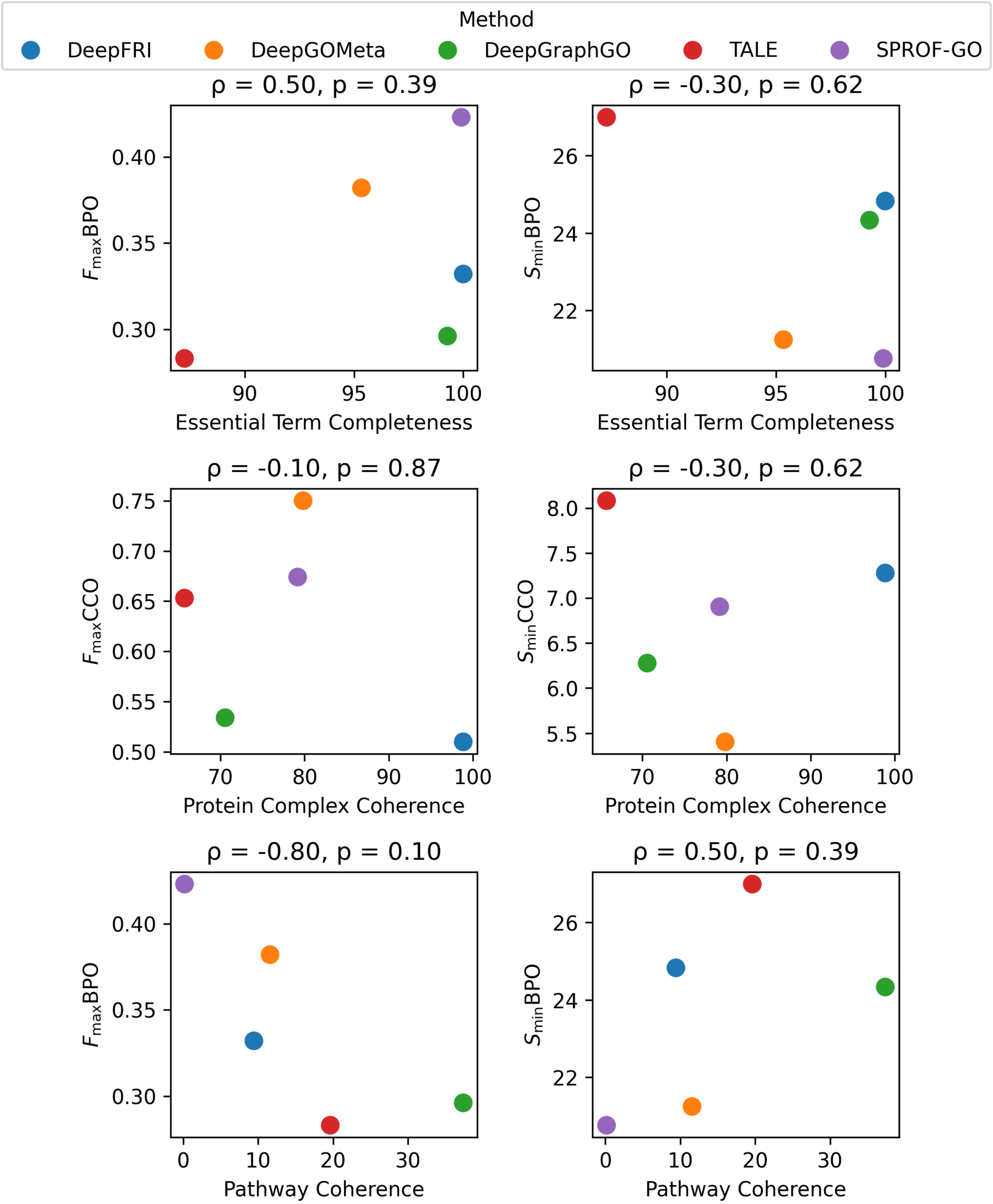


**Fig. S5.** Spearman correlations between CAFA evaluation metrics ($F_{\max}$, $S_{\min}$) and framework evaluation metrics (Essential term completeness, protein complex coherence, and pathway coherence).

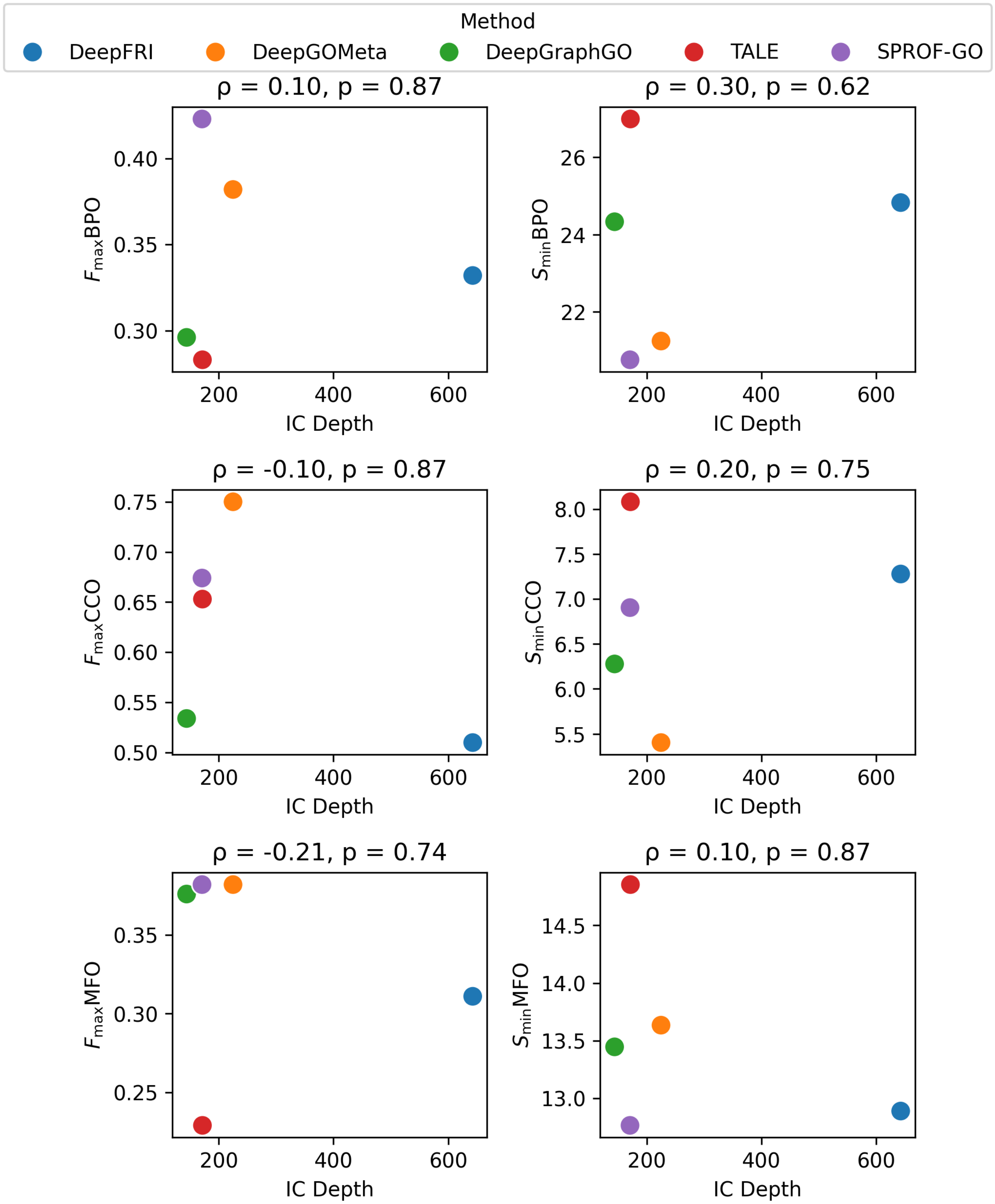


**Fig. S6.** Spearman correlations between evaluation metrics ($F_{\max}$, $S_{\min}$) and Information Content (IC) depth for all GO subdomains (BPO, CCO, and MFO) based on function prediction method results.

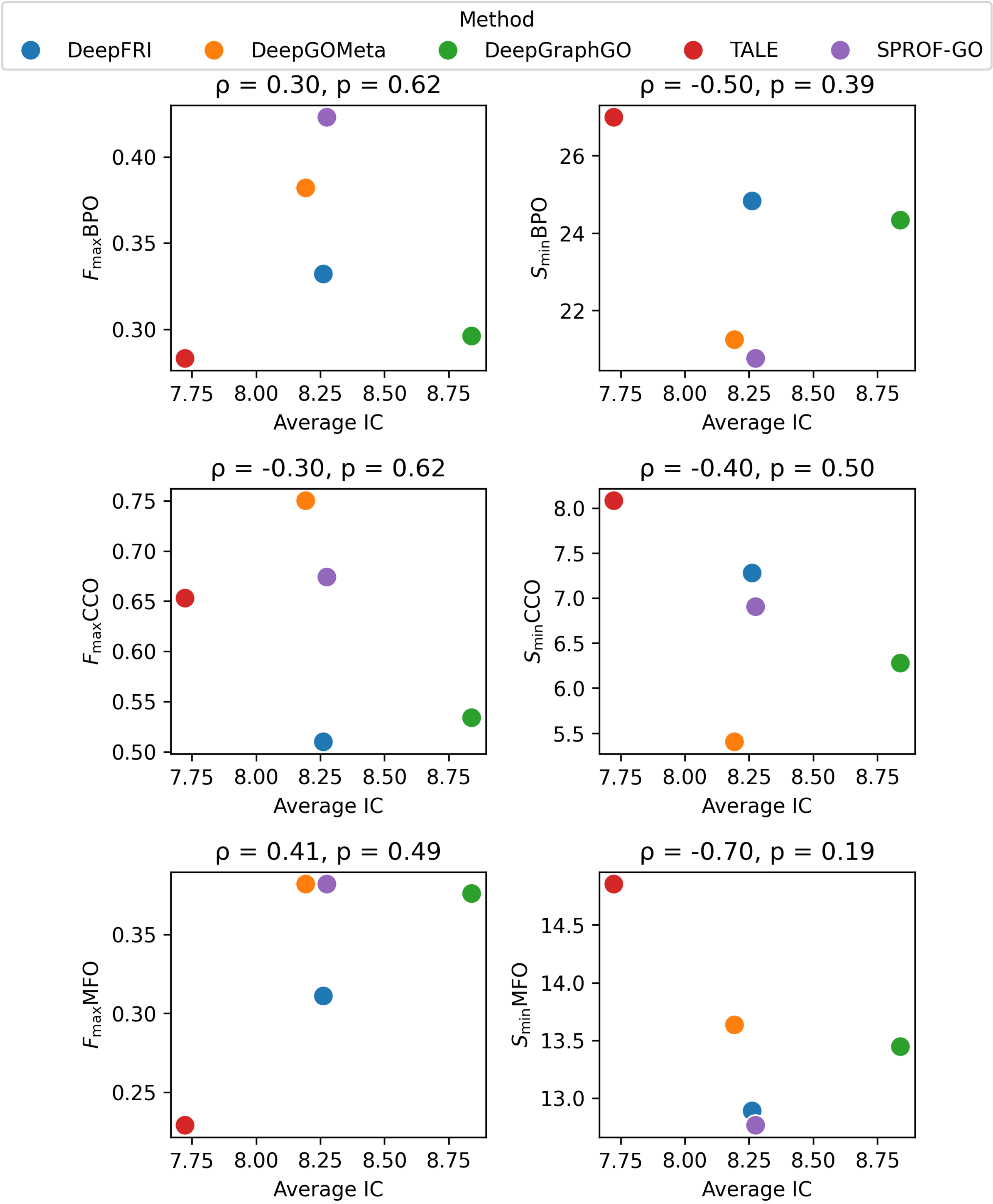


**Fig. S7.** Spearman correlations between CAFA evaluation metrics ($F_{\max}$, $S_{\min}$) and average Information Content (IC) for all GO subdomains (BPO, CCO, and MFO) based on function prediction method results.
